# DNA G-quadruplex is a transcriptional control device that regulates memory

**DOI:** 10.1101/2023.01.09.523337

**Authors:** Paul R. Marshall, Qiongyi Zhao, Joshua Davies, Wei-Siang Liau, Yujin Lee, Dean Basic, Ambika Periyakaruppiah, Esmi L. Zajaczkowski, Laura J. Leighton, Sachithrani U. Madugalle, Mason Musgrove, Marcin Kielar, Hao Gong, Haobin Ren, Lech Kaczmarczyk, Walker S. Jackson, Alon Chen, Robert C. Spitale, Timothy W. Bredy

## Abstract

The conformational state of DNA fine-tunes the transcriptional rate and abundance of RNA. Here we report that DNA G-quadruplex (G4-DNA) accumulates in neurons in an experience-dependent manner, and that this is required for the transient silencing and activation of genes that are critically involved in learning and memory. In addition, site-specific resolution of G4-DNA by dCas9-mediated deposition of the helicase DHX36 impairs fear extinction memory. Dynamic DNA structure states therefore represent a key molecular mechanism underlying memory consolidation.

**One-Sentence Summary:** G4-DNA is a molecular switch that enables the temporal regulation of the gene expression underlying the formation of fear extinction memory.

## Main Text

The extinction of conditioned fear is an evolutionarily conserved behavioral adaptation that is critical for survival. Like other forms of learning, long-lasting memory for fear extinction depends on coordinated changes in gene expression, particularly in the infralimbic region of the medial prefrontal cortex (ILPFC) (*1–3*). In recent years, we and others have shown that this process involves a tightly controlled interplay between the transcriptional machinery and epigenomic mechanisms (*4*). DNA is significantly more persistent than RNA, protein, or lipids in the cell, and the mechanisms surrounding its regulation are therefore key to understanding behavioral adaptation (*5*). Although DNA and histone modifications have long been associated with neuronal plasticity and memory (*6–12*), little is known about how local changes in DNA structure impact experience-dependent gene expression. This is because the relationship between DNA structure and function has primarily been attributed to the right-handed double helix, B-DNA, first described by Watson, Crick and Franklin. However, DNA can adopt more than 20 different conformational states, several of which have been linked to transcriptional activity. In addition, the biochemical conditions that promote dynamic conformational changes in DNA, including the influx of calcium, potassium, and sodium ions, are also directly involved in driving neuronal gene expression, suggesting that dynamic changes in DNA structure may be a critically important mechanism of memory.

We recently discovered that neurons can assume a left-handed conformational state (Z-DNA) in response to neural activity, which is critical for modulating the qualitative aspects of transcription and the strength of fear-related memories (*13*). However, whether other DNA structures also regulate gene expression essential for memory stability is completely unexplored. G-quadruplex DNA (G4-DNA), which accumulates when guanines fold into stable four-stranded DNA structures, is known to protect DNA during replication (*14*), is involved in class-switch recombination in immune cells (*15*), and dynamically regulates transcription in a variety of cell types (*16*). Like Z-DNA, G4-DNA is stabilized by ions that are highly abundant in activated neurons (*17*) and the folding kinetics of G4-DNA ranges from milliseconds to minutes, which overlaps temporally with the rate of transcription associated with learning. Moreover, G4-DNA helicases such as the DEAD-Box helicase 36 (DHX36) mediate the resolution of G4-DNA structure, a process that is strongly associated with the modification of transcription (*18*). We therefore posited that G4-DNA is involved in the regulation of experience-dependent gene expression and memory. We tested the hypothesis that the formation of G4-DNA and its resolution by DHX36 alter the rate of learning-induced transcription in the medial prefrontal cortex (mPFC), and that this process is directly related to the formation of fear-related memories.

### G4-DNA is regulated by DHX36 during fear and extinction learning

To determine if G4-DNA accumulates in response to fear extinction learning, we fear conditioned 10-16 week old C57 mice using a standard tone-shock pairing protocol. We then exposed these mice in a novel context to either a 10, 30, or 60 conditioned stimulus (CS) extinction (EXT) protocol, or to an equivalent time without re-exposure to the previously conditioned stimulus (retention control (RC)) and extracted the mPFC after both tasks. An examination of global levels of G4-DNA and mRNA expression of the G4-DNA helicase DHX36 revealed an increase in G4-DNA, 1h post-fear training (fig. S1A), which was followed by an increase in DHX36 mRNA 5h post-fear training (fig. S1B) and at 10 CS EXT (Fig. 1A). This suggested an effect of DHX36 on G4-DNA during the late phase of fear consolidation and during extinction learning. G4-DNA sequencing (G4-seq (*19*)) on DNA derived from mPFC neurons during the early phase of extinction learning (10 CS EXT and RC) revealed that 10 CS EXT training led to an increase in the percentage of G4-DNA reads 5 kilobases (kb) upstream or downstream of the transcription start site (proximal TSS) and within introns (Fig. 1B).

**Fig 1.**
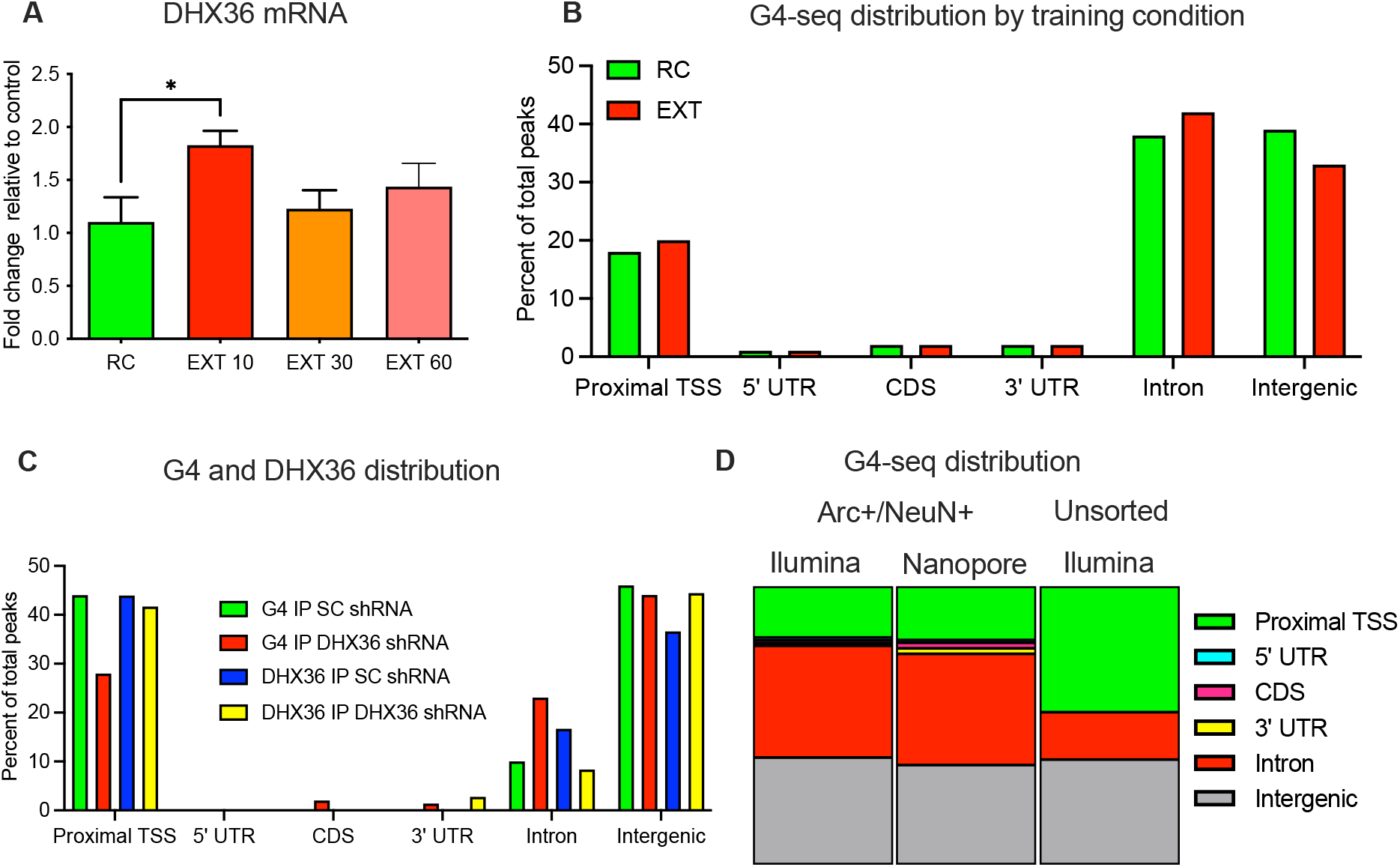
Neuronal genome-wide distribution of G4-DNA and DHX36 is influenced by learning and depends on cell-type and activation state. **(A)** DHX36 mRNA expression is transiently induced by fear extinction learning (EXT 10 CS (F_3,15_ = 2.882, p = 0.07; Dunnett’s post-hoc test; RC vs EXT 10, *p<0.05). Ctx A- Context A; FC-fear conditioning; EXT-extinction trained; RC-retention control). **(B)** Distribution of G4-DNA from RC and EXT trained mice. 5’ untranslated region (5’ UTR), proximal TSS (5kb +/− transcription start site), CDS (coding region), 3’ untranslated region (3’ UTR). **(C)** Distribution of G4-DNA and DHX36 in EXT trained animals treated with scrambled control virus (SC) and DHX36 shRNA virus **(D)** Distribution of G4-DNA in activated and unsorted neurons on short (Illumina) and long-read (nanopore) sequencing platforms.

To identify G4-DNA regions that were actively regulated during extinction we designed an shRNA against DHX36, which produced a significant reduction in DHX36 mRNA (fig. S1C) and increased G4-DNA (fig. S1D). Mice were fear conditioned and infused with scrambled control (SC) shRNA or DHX36 shRNA into the ILPFC followed by extinction training. We then performed G4-DNA-seq and DHX36 immunoprecipitation-seq (Data S1). The most significant change in distribution occurred within introns, with DHX36 shRNA increasing the accumulation of G4-DNA and decreasing DHX36 occupancy within these genomic regions (Fig. 1C). To further identify G4-DNA regulatory sites in neurons we performed G4-DNA-immunoprecipitation, short and long-read sequencing on neurons that had been selectively activated by learning (Data S2). A direct comparison was made between cells identified as activated neurons based on neuronal nuclei (NeuN) and activity-regulated cytoskeleton (Arc) expression (Arc+/NeuN+ G4-DNA), and unsorted homogenates (with a high proportion of Arc-/NeuN+ neurons). Although the distribution of reads was nearly identical for long-read and short-read sequencing, we observed a greater proportion of reads in introns over the proximal TSS in the activated neuron population (Fig. 1D). Together, these data suggest that neuronal activity drives a conformational shift in G4-DNA at the TSS toward the accumulation of G4-DNA within intronic regions.

We next overlapped these data sets in the USC genome browser and the integrated genomics viewer (IGV) to reveal common targets, which had 1) G4-DNA in all three G4-DNA seq experiments, 2) an increase in G4-DNA when comparing DHX36 shRNA to control, and 3) DHX36 occupancy and a reduction in DHX36 following knockdown. We selected eight targets for validation based on their known association with neuronal plasticity: ying-yang 1 (yy1), nesprin 2 (syne2), adducin 1 (add1), regulating synaptic membrane exocytosis 4 (Rims4), potassium large conductance calcium-activated channel, subfamily M, beta member 4 (Kcnmb4), RPTOR independent companion of MTOR, complex 2 (Rictor), gephyrin (Gphn) and neural cell adhesion molecule L1-like (Chl1).

Chl1 and Gphn exhibited the most robust G4-DNA signal in introns, with large differences in G4-DNA and DHX36 signal being observed between the two conditions (fig. S1E). Validation with G4-DNA IP in combination with formaldehyde-assisted isolation of regulatory elements (FAIRE-Seq) revealed an increase in G4-DNA in both genes following the reduction in DHX36 (fig. S1, F and G). Although G4-DNA and DHX36 binding increased in Rims4, Kcnb4, and Rictor (fig. S2, A to I) during extinction learning, Yy1, Syne2, and Add1 showed the opposite effect, a finding that was most evident in Arc+ activated neurons (fig. S3, A to I). We therefore concluded that G4-DNA is regulated by DHX36 in neurons at genes that are of critical importance for neuronal plasticity, and that this occurs in an experience-dependent manner.

### DHX36 knockdown transiently increases mRNA expression during extinction learning

It is well established that an increase in G4-DNA, either by stabilizing compounds or by G4-DNA helicases such as DHX36, leads to reduced transcription (*16*). This is thought to occur as a result of G4-DNA impeding the progression of RNA polymerase II (Pol II). In contrast, G4-DNA has also been shown to enhance transcription when it forms proximal to the TSS, which may occur because of G4-DNA-mediated maintenance of the transcription bubble, which then primes subsequent gene expression (*16*). To determine which of the two primary models accounts for the G4-DNA-mediated effects on experience-dependent transcriptional activity in neurons, we co-infused a self-deleting Cre recombinase and either SC shRNA or DHX36 shRNA into the ILPFC of mice expressing a Cre-activated UPRT cassette (*20*). To label nascent RNA, 4-thiouracil (4-TU) was infused 2 hours prior to exposure to a 30CS EXT protocol, followed by sulfide-mediated click-chemistry enrichment to capture nascent RNAs expressed in response to extinction learning followed by sequencing (Fig. 2A).

**Fig 2.**
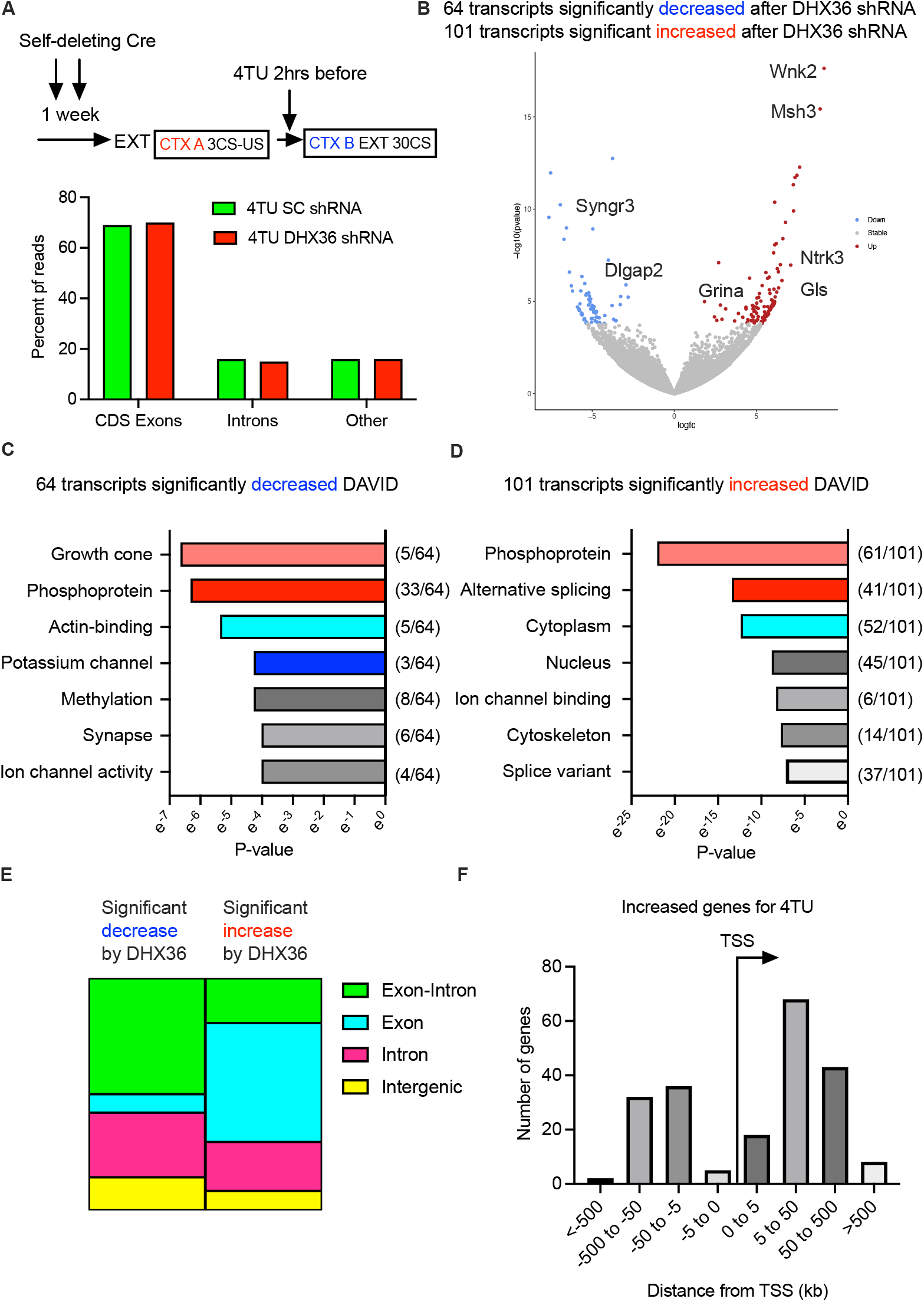
Experience-dependent nascent RNA expression is modulated by G4-DNA. **(A)** Timeline of training and 4-Thiouracil (4TU) infusions, and distribution of reads across the transcriptome **(B)** Volcano plot of 4TU labelled transcripts which significantly increased and decreased relative to control when DHX36 shRNA was infused **(C)** Database for Annotation, Visualization and Integrated Discovery (DAVID) plot comparing category to P-value for of all the significantly decreased transcripts **(D)** and all of the significantly increased transcripts **(E)** A visual representation indicating the percentage of 4TU reads from the significantly decreased and increased transcripts which occurred across introns and exons (Exon-Intron), only over exons (Exon), only over introns (Intron), or only over intragenic regions (Intragenic) **(F)** The number of significantly increased 4TU transcripts and their distances in kilobases relative to the closest annotated transcription start site.

A comparison between nascent RNA derived from SC- and DHX36 shRNA-treated mice revealed reduced expression in 64 genes following DHX36 knockdown, whereas 101 transcripts were found to increase (Fig. 2B and Data S4). Gene ontology analysis revealed that transcripts related to neuronal processes such as ion channels and actin binding were reduced (Fig. 2C). In contrast, increased expression was found in genes associated with alternative splicing as well as genes that are known to regulate ion channel binding (Fig. 2D). Further, decreased RNA expression mostly occurred across larger sections of the gene as indicated by a higher percent of significant reads in exon-intron spanning regions, whereas the increase in nascent RNA expression was detected primarily over exons (Fig. 2E). To determine whether the effect on transcription was the result of G4-DNA stabilization of the transcription bubble around the TSS, we analyzed the overlap between the regions of increased transcription, G4-DNA sites, and TSS (identified by altTSS), revealing a bias toward increases upstream rather than downstream of the TSS (Fig. 2F). These data suggest that, in the context of fear extinction learning, the accumulation of G4-DNA is associated with both reduced and increased nascent RNA expression, and that this occurs along a very short-term temporal scale.

### G4-DNA promotes polymerase II stalling within learning-related genes

Because nascent RNA-seq using metabolic labeling with 4-TU is more likely to detect regulatory mechanisms associated with enhanced transcription by sampling nascent RNA expression in a narrow temporal window, we next wanted to assess potential polymerase stalling related to G4-DNA transcriptional regulation. A separate cohort of animals was infused with SC and DHX36 shRNA followed by a 30CS EXT protocol. The mPFC was extracted post-training and subjected to both total RNA-seq and native elongation transcript sequencing (NET-seq), which enabled the quantification of Pol II-associated RNA and a comparison between total RNA and the binding of transcripts to Pol II in the same animal (*21*). Overall, we observed a significant reduction in transcripts following DHX36 shRNA treatment when sampling total RNA (Fig. 3A and Data S3). Although most transcripts exhibited no change in RNA levels or Pol II occupancy (Fig. 3B), 27% of genes exhibited a decrease in RNA expression (in the total RNA fraction) with a concomitant increase in Pol II binding (in the Pol II fraction), suggesting that Pol II stalling had occured in genes where G4-DNA accumulated but was not removed In fact, we observed a significant decrease in total RNA and a significant increase in the fraction of RNA associated with Pol II within all 8 of the validated G4-DNA targets (fig. S4, B to Q and Data S5). Further, in comparison to nascent RNA-seq, the few transcripts that increased following DHX36 shRNA treatment were more likely to be introns (Fig. 3C). However, there was a bias for the increase to occur upstream of the TSS, again supporting the interpretation that increased RNA expression may be governed by G4-DNA maintenance of the transcription bubble. Overall, these results indicate that stable G4-DNA sites primarily act as gene silencers in neurons.

**Fig 3.**
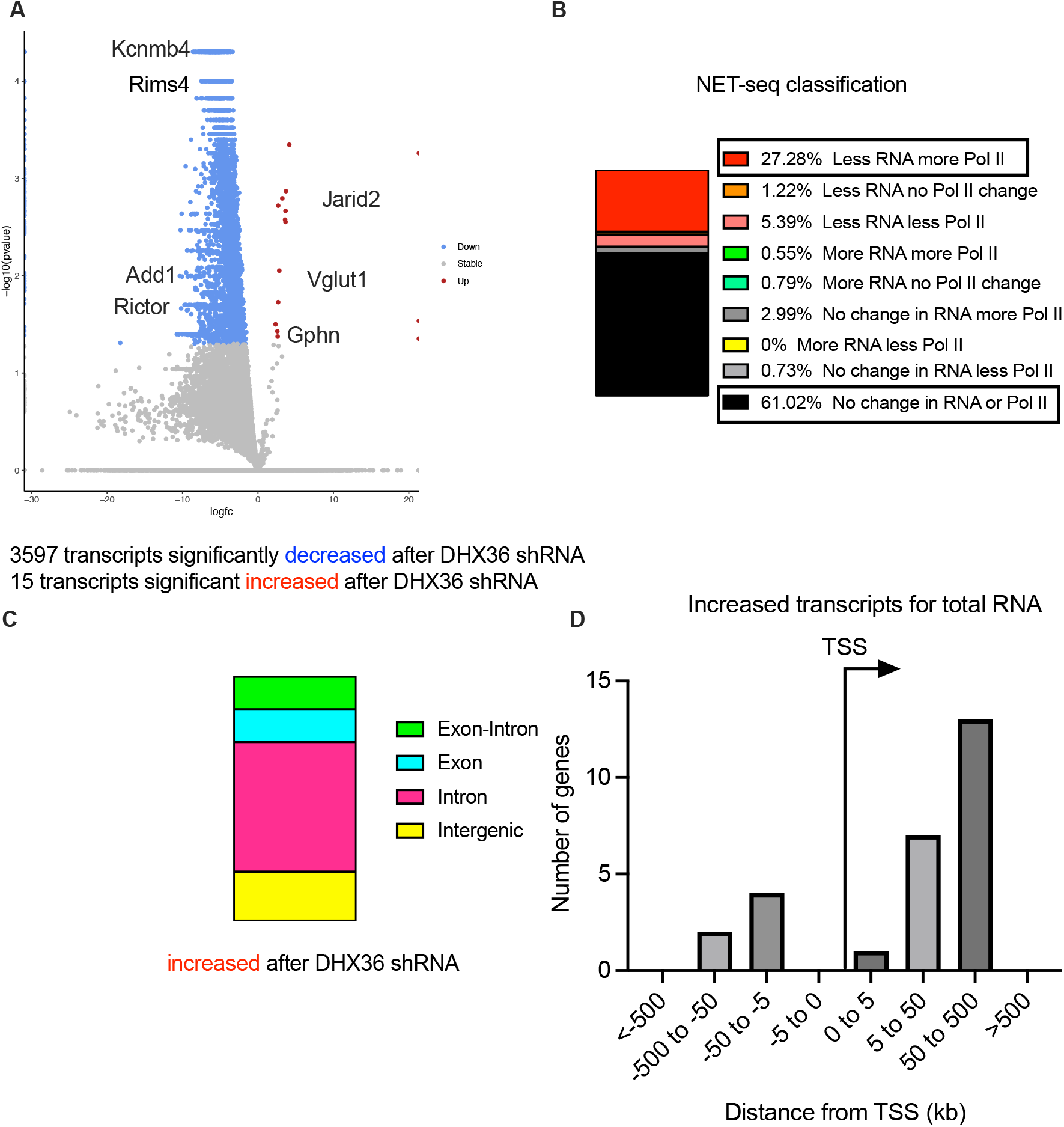
Pol II associated RNA is influenced by G4-DNA following DHX36 knockdown. **(A)** Volcano plot of total RNA expression following infusion of DHX36 shRNA relative to control **(B)** Plot comparing changes in total RNA levels to changes in the amount of Pol II bound RNA sorted by categories of increasing, decreasing or no change when comparing control virus to DHX36 shRNA infusion **(C)** A visual representation indicating the percentage of reads from the significantly increased total transcripts which occurred across introns and exons (Exon-Intron), only over exons (Exon), only over introns (Intron), or only over intragenic regions (Intragenic) **(D)** The number of significantly increased total transcripts and their distances in kilobases relative to the closest annotated transcription start site.

### Novel targets for G4-DNA-mediated transcriptional regulation

Our transcriptome-wide data provided support for both models of transcriptional regulation by G4-DNA. To further explore the role of G4-DNA in experience-dependent gene regulation, we next used a gene analysis of changes in RNA expression. A manual re-analysis of every significantly altered gene in the transcriptome-wide data, by inputting each coordinate into the USC genome browser and overlapping it with the G4-DNA peak data, as well as our own and previously published datasets, revealed many other examples of this phenomenon (fig. S6). The RNA products overlapping with G4-DNA did not appear to share a common subtype, as some occurred over annotated and putative long noncoding RNAs as well as microRNAs, whereas the Gphn locus appeared to encode a circular RNA as evidenced by its resistance to RNAse R treatment (fig. S6). Together, these findings suggest that although increased RNA expression can be explained by G4-DNA being maintained proximal to the TSS, and decreased RNA expression is associated with Pol II stalling, both short- and long-term changes in G4-DNA are also associated with a novel subclass of noncoding RNAs, the functional relevance of which remains to be investigated.

### DHX36 is required for the consolidation of fear extinction memory

Having established that G4-DNA has a significant impact on transcription we next sought to investigate its effect on learning and memory, *in vivo*. Specifically, to determine if G4-DNA regulates memory stability, DHX36 shRNA was infused into the ILPFC of mice immediately after fear training, followed by fear extinction training a week later and two tests in both the original context (A) and extinction context (B) (Fig. 4A). DHX36 shRNA had no effect on within-session extinction learning (fig. S7A); however, extinction memory was impaired in both contexts (Fig. 4B-C). A separate cohort of animals was infused with SC and DHX36 shRNA into the prelimbic region of the mPFC (Fig. 3D), which is known to regulate the acquisition and consolidation of fear memory. Curiously, DHX36 shRNA led to a significant impairment in the acquisition of cued fear (Fig. 4E) but had no significant effect on expression at test (Fig. 4F). Together these data suggest that preventing the removal of G4-DNA impairs memory processes.

**Fig 4.**
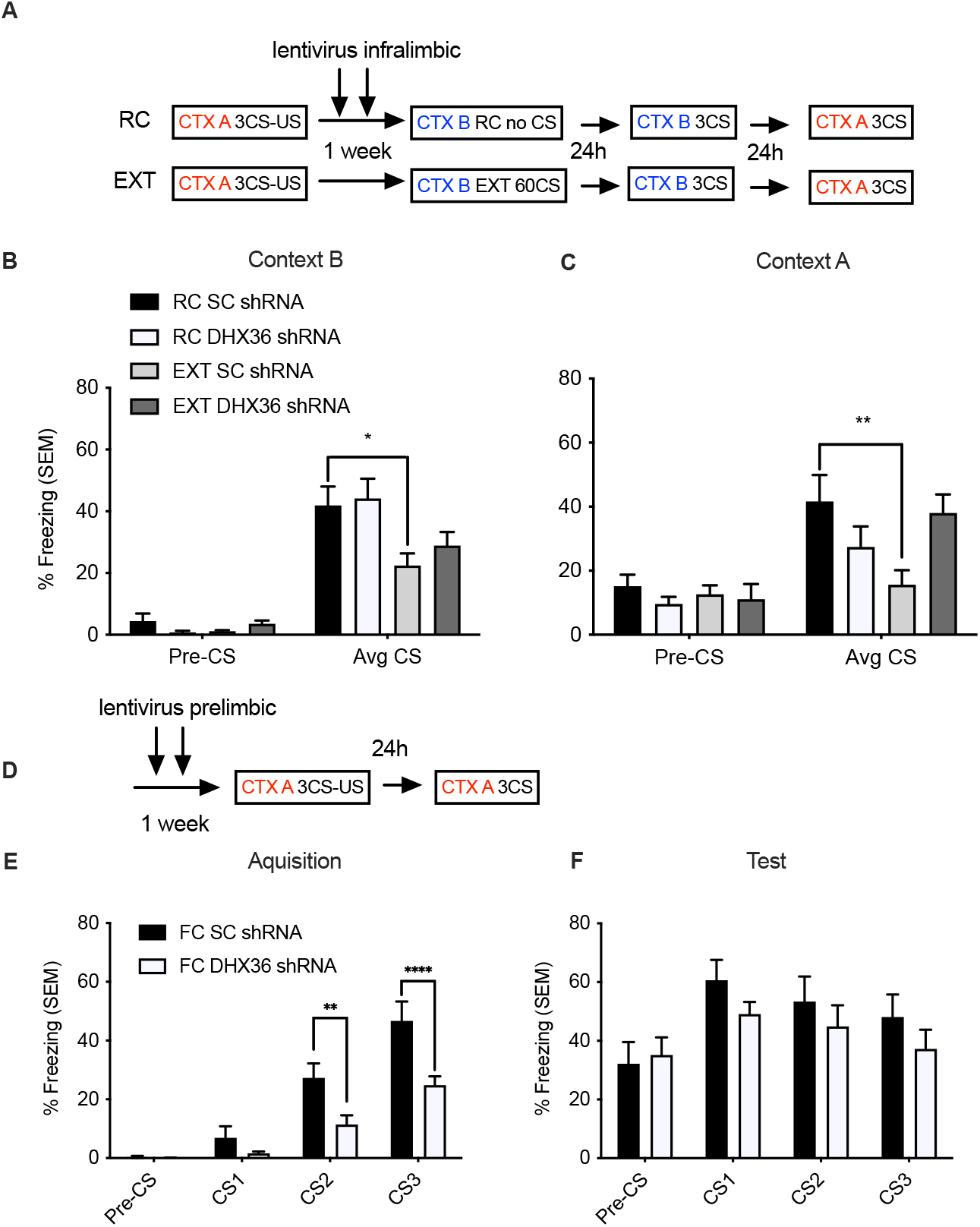
G4-DNA effects both fear extinction and fear acquisition. **(A)** Timeline of training **(B)** Infusion of DHX36 shRNA into the infralimbic PFC led to a significant impairment in memory for fear extinction following a 60 CS training protocol in context B (n=8/group, one-way ANOVA F_3,39_= 4.466, p<0.01, Dunnett’s posthoc test; RC Scrambled vs. EXT Scrambled, *p<0.05). EXT-extinction trained; RC-retention control; Avg CS-Average of 3 conditioned stimulus exposures). **(C)** DHX36 shRNA also led to a significant impairment in memory for fear extinction following a 60 CS training protocol in context A (n=8/group, two-way ANOVA F_1,39_= 4.215, p<0.05, Dunnett’s posthoc test; RC Scrambled vs. EXT Scrambled, **p<0.01). **(D)** Infusion of DHX36 shRNA into the prelimbic PFC led to a significant impairment in memory for acquisition (F_3,56_= 4.335, p<0.01, Sidak posthoc test; FC Scrambled vs. FC DHX36, **p<0.01 and ****p<0.0001)

### Targeted reduction in G4-DNA regulates fear memory, depending on the gene target

One limitation of manipulating DHX36 is that it targets G4-RNA as well as G4-DNA. To overcome this issue and more directly define a role for DHX36 in the regulation of G4-DNA, we designed a dcas9-DHX36 construct, which can be directed to specific genomic loci to drive the resolution of G4-DNA. We selected Gphn and Chl1 based on their known roles in plasticity and memory and our observation of a learning-induced role for G4-DNA-mediated effects on gene expression at these loci. We found no effect of dcas9-DHX36 at the Gphn locus on within-session extinction (fig. S7B); however, extinction memory was significantly impaired (Fig. 5C). In contrast, although there was no effect on within-session extinction when Chl1 was targeted by dcas9-DHX36, a significant reduction in fear expression in the Chl1-RC group and a significant increase in transcription at this G4-DNA site were observed (Fig. 5E). We then performed a second experiment where dcas9-DHX36 was infused prior to fear learning. Although targeting DHX36 to the Chl1 locus led to a modest effect on fear during acquisition and retrieval, the same manipulation at the Gphn locus produced significant impairments in both the acquisition and expression of cued fear (fig. S7 D-E). In addition, we validated that the dcas9-DHX36 significantly reduced G4 levels at the Gphn and Chl1 locus (fig. S7 F-G). Together these data confirm that G4-DNA serves to regulate learning-induced gene expression in a state-dependent and gene-specific manner, which can have opposing effects depending on the phase of memory formation.

**Fig 5.**
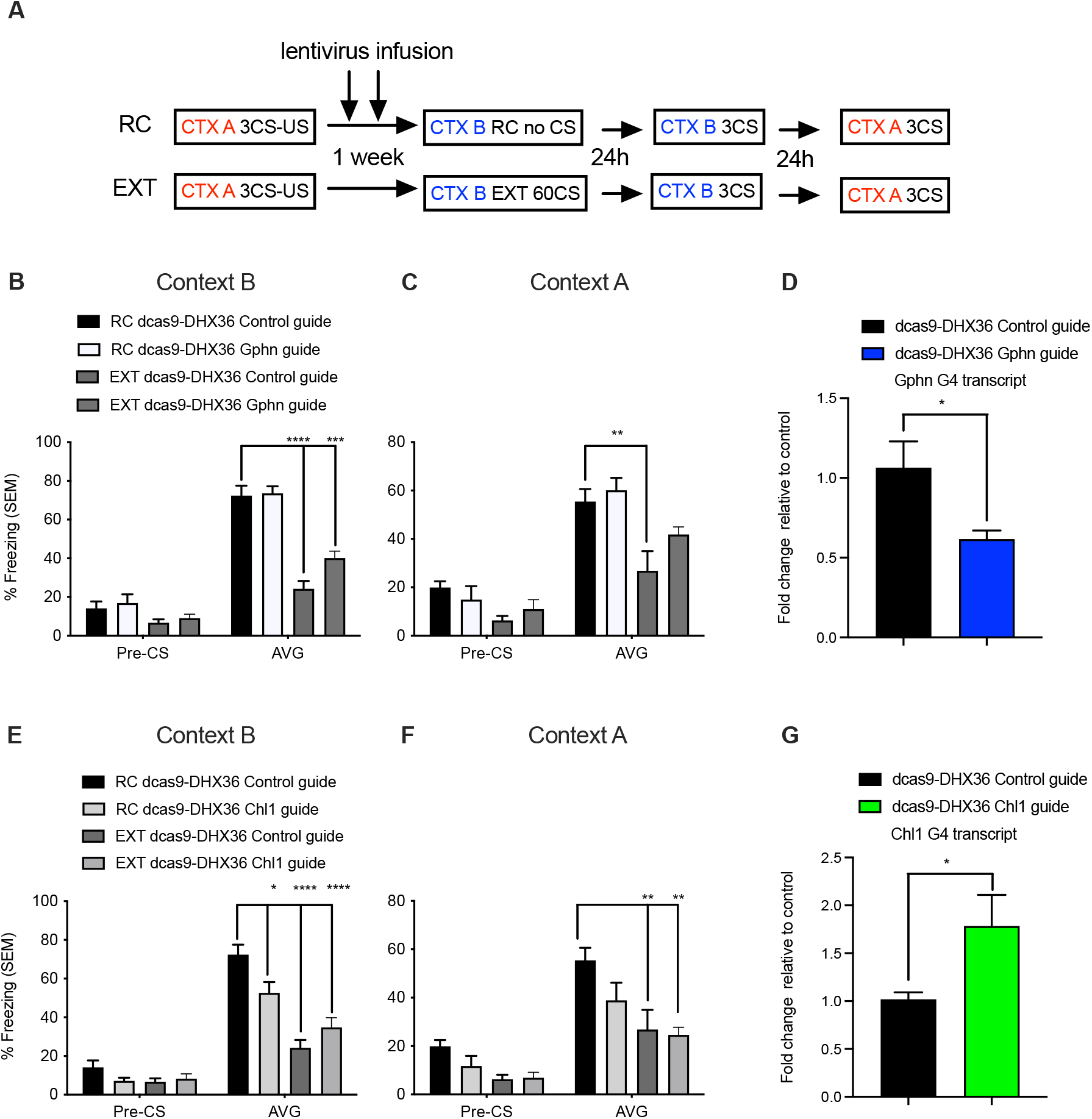
Targeted resolution of G4-DNA at Chl1 and Gph impairs the consolidation of fear-memory. **(A)** Timeline of training **(B)** Lentivirus infusion into the infralimbic PFC of a CRISPR-dCAS9-DHX36 construct directed to either a control or Gphn locus led to a significant impairment in memory for fear extinction following a 60 CS training protocol in context B (**(C)** as well as context A ((**D)** CRISPR-dCAS9-DHX36 directed to the Gphn locus leads to a significant decrease in mRNA expression (n=7/group, t-test t_11_ = 2.76, *p<0.05). **(E) (F) (G)** CRISPR-dCAS9-DHX36 directed to the Chl1 locus led to a significant increase in mRNA expression (n=6/group, t-test t_11_= 2.48, *p<0.05).

## Discussion

The accumulation of G4-DNA has classically been associated with telomere maintenance (*14*) and, more recently, with class-switch recombination in immune cells (*15*). Although it has been previously observed in neurons, it was thought to reflect genome instability and DNA damage (*22*) and autophagy (*23*, *24*). Here we have found that G4-DNA is regulated by DHX36 in genes with known roles in neuronal plasticity, with the most pronounced effects occurring in introns. Thus, although persistent G4-DNA may produce damage and impaired transcription in some cell types, it is also involved in neural plasticity and appears to be temporally regulated by DHX36, implying a role for G4-DNA-specific binding proteins and helicases in neurons (*25*, *26*). In this respect, DHX36 regulates G4-DNA primarily in the TSS and introns in neurons, whereas in immune cells BMI1 promotes the accumulation of G4-DNA and subsequent regulation of L1-containing transcripts, suggesting that different G4-DNA-related binding proteins may regulate different subregions of the genome (*27*).

G4-DNA has been linked to decreased RNA expression by disrupting Pol II readthrough (*28*). We also observed robust decreases in total RNA, which were accompanied by the accumulation of Pol II following an increase in G4-DNA by DHX36 knockdown. This provides further support for the effect of Pol II RNA stalling on RNA expression and extends this to include neurons involved in fear and extinction learning. We also found that G4-DNA promotes the expression of specific transcripts both downstream of G4-DNA and directly at this site. These observations, in combination with the changes in G4-DNA induced by DHX36 at different timepoints for different genes, implies an activity-regulated switch such that G4-DNA, if stabilized, can inhibit transcription on a long-term timescale, followed by rapid initiation of transcription following targeted release of G4-DNA. This is supported by our data on the Gphn gene locus whereby extended G4-DNA following DHX36 knockdown resulted in a global reduction of Gphn, that robust neuronal activity increased G4-DNA and reduced RNA expression with subsequent behavioral training or weak depolarization, leading to G4-DNA release and the triggering of transcription. These data further highlight the need to better understand the mechanisms underlying G4-DNA formation, stability, and resolution across a variety of cell types and activity states.

It is evident that G4-DNA regulation is required for memory processes. Although we observed no effect on fear extinction learning when G4-DNA was increased in the ILPFC, the consolidation of fear extinction memory was impaired. One caveat is that the manipulation of DHX36 can also influence G4-RNA. Therefore, using the dCas9 system, we manipulated G4-DNA at the Gphn and Chl1 loci, respectively, finding that both led to significant changes in RNA expression associated with the G4-DNA and influenced learning and memory. Target resolution of G4-DNA at the Chl1 locus led to impaired fear memory, whereas the same manipulation at the Gphn locus impaired fear extinction memory. Subsequent manipulations prior to the acquisition of fear further confirmed that a DHX36-mediated reduction in G4-DNA at the Gphn locus directly affects fear acquisition. These findings suggest a switch-like influence of G4-DNA on transcription as either chronically stabilizing G4-DNA by reducing DHX36, or constitutively reducing G4-DNA by driving DHX36 to specific sites along the genome, leads to overall impairments in memory.

In summary, we have discovered that G4-DNA is directly involved in modulating fear-related memory and that the global model for the regulation of transcription by G4-DNA is dependent on both the cell type and its activation state. Historically, the accumulation of G4-DNA in neurons has been thought to reflect DNA damage and impaired transcriptional activity; however, when G4-DNA is temporally restricted by DHX36, it clearly plays a permissive role in experience-dependent neuronal plasticity. G4-DNA is therefore a critical molecular switch underlying the regulation of neuronal transcription and the consolidation of memory.

## Author contributions

Conceptualization: PRM & TWB
Investigation: PRM, JD, YL, AP, ELZ, LJL, SUM, MM, MK, HG, HR
4TU-seq and NET-seq: PRM, RCS & TWB
Tagger Experiments: LK, WJ
Bioinformatics: QZ, PRM, DB
Plasmid Design: WSL, AC
Funding acquisition: PRM & TWB
Project administration: TWB
Supervision: TWB
Writing – original draft: PRM & TWB
Writing – review & editing: PRM & TWB

## Data and materials availability

All data are available in the main text or the supplementary materials.

## Supplementary Materials

S1. Validation of DHX36 shRNA

S2. DHX36 and G4-DNA time course for increased G4-DNA targets

S3. DHX36 and G4-DNA time course for decreased G4-DNA targets

S4. Polymerase II stalling and reduced RNA at validated targets

S5. Chl1 and Gphn total and nascent RNA overlap with G4-sites

S6. Sites of G4-DNA and significant RNA change across the transcriptome

S7. Fear and extinction behaviour during viral manipulation

