## Supplementary Materials for "DNA G-quadruplex is a transcriptional control device that regulates memory"

**This PDF file includes:**

Materials and Methods
Supplementary Text
Figs. S1 to S7
Captions for Data S1 to S8

**Materials and Methods**

Animals 9-11 week-old C57BL/6 male mice were housed four per cage, maintained on a 12h light/dark schedule, and allowed free access to food and water. To allow for identification of behavioural outliers, animals were transferred to pair-housed conditions and split with a plexiglass divider at least 24h prior to training. All testing was conducted during the light phase in red-light-illuminated testing rooms. All animal use and training, including the use of embryos, followed protocols approved by the Animal Ethics Committee of the University of Queensland and in accordance with the Australian Code for the Care and Use of Animals for Scientific Purposes (8th edition, revised 2013).

Lentiviral delivery and behavioural analysis Double cannulae (PlasticsOne) were implanted in the anterior posterior plane, along the midline, into the infralimbic prefrontal cortex (ILPFC). The injection locations were centred at +1.8 mm in the anterior-posterior plane (AP), and -2.8 mm in the dorsal-ventral plane (DV). For prelimbic prefrontal cortex the injection locations were centred at +1.8 mm in the anterior-posterior plane (AP), and -1.8 mm in the dorsal-ventral plane (DV). Animals were then separated into single housing and given at least one week to recover before being behaviourally trained. Following the first day of training a total volume of two microliters of lentivirus per hemisphere was infused via two one microliter injections over a 48h period. Mice were first fear conditioned, followed by 2 lentiviral infusions 24 hours post-fear conditioning, and, after a one-week of incubation, the mice were either extinction trained or just exposed to context B for an equivalent period of time. In brief, this training consisted of two contexts (A and B). Both conditioning chambers (Coulbourn Instruments) had two transparent walls and two stainless steel walls with a steel grid floors (3.2 mm in diameter, 8 mm centers); however, the grid floors in context B were covered by a flat white plastic transparent surface. Context A was also sprayed with a dilute lemon smell, and context B was sprayed with a dilute vinegar smell to minimize context generalization. Cameras within the boxes captured movement and were processed automatically with a freezing measurement program (FreezeFrame). The training protocol consisted of 120 s pre-fear-conditioning incubation, followed by three pairings of a 120 s, 80dB, 16,000 Hz tone (CS) co-terminating with a 1 sec (2 min intervals), 0.7 mA foot shock, or unconditioned stimulus (US). Mice were matched into equivalent treatment groups based on freezing during the third training CS. For extinction, mice were exposed in context B in which they habituated to the chamber for 2 min, after which the extinction training (EXT) comprised 10, 30 or 60 non-reinforced 120 s CS presentations (5-s intervals). For the retention control (RC) animals, context exposure was performed following fear conditioning, but without presentation of the tones. For the retention tests, all mice were returned to either context A or B and following a 2 min acclimation (used to minimize context generalization), freezing was assessed during three 120s CS presentations (120 s intertrial interval). Memory was inferred by the percentage of time spent freezing during the tests.

mPFC extractions and tissue preparation

Following the ending of behavioural training animals were transported to a separate room. Here cervical dislocation was performed followed by immediate extraction of the medial prefrontal cortex on ice. Tissue was then dounced in 2ml tissue grinder (Kimble Chase) with buffers and inhibitors related to downstream procedures.

Primary cortical neurons Cortical tissue was isolated from embryonic day 16 embryos. Primary cortical neurons were isolated by removing the skull and meninges with fine-tipped tweezers. Cells were then mixed into a medium solution comprising Neurobasal medium (GIBCO 21103) containing 5% serum, B27 supplement (GIBCO 17504-044) and 0.5-1% Penicillin-Streptomycin (GIBCO 15140), made homogenous with gentle pipetting. The cells were then passed through a 40 micrometer cell strainer (BD Falcon 352340) and plated onto 6-well cell culture dishes coated with poly-L- ornithine (Sigma P2533) at a density of 1 million cells per well.

RNA and DNA extraction Both cultured cells and tissue from mice were extracted and then placed in PBS. Gentle pipetting was used for *in vitro* preparations to generate a homogenous solution. Tissue was prepared by dounce homogenization in 500 µl of PBS and RNA/DNA was extracted. For RNA, the Trizol reagent was used according to the manufacturer's instructions (Invitrogen). DNA extraction was carried out using the DNeasy Blood & Tissue Kit (Qiagen) with RNAse A (5 prime), RNAse H and RNAse T1 treatment (Invitrogen). Both extraction protocols were conducted according to the manufacturer's instructions. The concentration of DNA or RNA was measured by Qubit assay (Invitrogen).

qRT-PCR 1μg of RNA was used for cDNA synthesis prepared from the Quantitect Reverse Transcription Kit according to the manufacture's protocol (Qiagen). Quantitative PCR was then performed on a RotorGeneQ (Qiagen) real-time PCR cycler with SYBRGreen Master mix (Qiagen), using primers for target genes and beta actin or phosphoglycerate kinase as an internal control. The threshold cycle for each sample was chosen from the linear range and converted to a starting quantity by interpolation from a standard curve run on the same plate for each set of primers. All mRNA levels were normalized for each well relative to the internal control using the ΔΔCT method, and each PCR reaction was run in duplicate for each sample and repeated at least twice.

DHX36 and DHX36 scrambled control (SC) knockdown lentiviral constructs Lentiviral plasmids were generated by inserting either DHX36 or SC shRNA using the following sequences or DHX36: GATCCCCGCCATCTAG CTACTATAAATTCAAGAGATTTATAGTAGCTAG ATGGCTTTTTC DHX36 SC: GATCCCCAGTTCATTAGGCTAACGTATTTCAAG AGAATACGTTAGCCTAATGAACTTTTTTTC immediately downstream of the human H1 promoter in a modified FG12 vector (FG12H1, derived from the FG12 vector originally provided by David Baltimore, CalTech). Lentivirus was prepared and maintained according to previously published protocols (*1*).

DHX36-dcas9 constructs Murine DHX36 was generated by adding XhoI and MfeI restriction enzyme sites to commercially available DHX36 cDNA clone (MBS1278832 MyBioSource). The cDNA was cloned into the Syn1-dcas9 vector (Addgene 114194) and guide RNA into mCherry vector (Addgene 114199). All plasmids were sequence verified by sanger sequencing.

FACS The procedure for sorting activated neurons for cDNA preparation and ChIP-seq was modified from published protocols(*2*). Briefly, following sample preparation for FACS the identified population of neurons, as indicated by their high intensity in the 488nm channel, was further split into two populations which had intensity in the 647nm Arc channel above the upper part of the non-NeuN population (high Arc) or below it (low Arc; see Figure 1b). 250,000 cells from each sample was then taken and four biological replicates were pooled to make one for further processing to reach at least the 1 million cells required for reliable chromatin immunoprecipitation.

Chromatin and carrier ChIP immunoprecipitation (ChIP) Standard ChIP was performed following modification of the Invitrogen ChIP kit protocol. Lysate from cells or tissue was fixed in 1% formaldehyde and cross-linked cell lysates were sheared by Covaris in 1% SDS lysis buffer to generate chromatin fragments with an average length of 300bp. For samples not being subjected to sequencing, 1 million yeast (kindly provided by Micheal Kobor) per sample were spiked in prior to fixation to enhance antibody-target interactions when the cell count was low (*3*). Following shearing, the chromatin was then immunoprecipitated following previously published protocols for G4 ChIP-seq (*4*), using the G4-DNA antibody (MABE917 Sigma-Aldrich) and Anti-FLAG M2 Beads (M8823-1ML Sigma Aldrich). Other ChIPs were carried out with, 5hmC (supplier), anti-RNA polymerase II CTD repeat YSPTSPS (phospho S5) antibody (ab5131), anti-POLR3A antibody (ab96328), or normal rabbit IgG (Santa Cruz), overnight at 4°C. Protein-DNA-antibody complexes were then precipitated with Dynabeads Protein G (Thermofisher 10003D) or Dynabeads MyOne Streptavidin C1 (Thermofisher 65001), for 1h at 4°C, followed by three washes in low salt buffer, and three washes in high salt buffer for protein G beads, and 3 washes in biotin wash buffer, followed by 2 washes in PBS. The precipitated protein-DNA complexes were eluted from the antibody with 1% SDS and 0.1M $NaHCO_3$ and incubated for 4h at 65C in proteinase K. Following proteinase K digestion, phenol-chloroform extraction, and ethanol precipitation, the samples were subjected to qPCR using primers specific for 200bp segments corresponding to the target regions. For FAIRE-seq experiments the lysate was split into two equal volumes and half was treated with the standard protocol as described above. The other half followed the same protocol but during the Proteinase K step no enzyme was added.

ChIP-Seq data analysis Bioinformatics analysis was performed as previously reported(*2*). After removing the duplicate reads, low mapping quality reads and paired-end reads that were not properly aligned, MACS2 (v.2.1.1.20160309) were used to call peaks for each sample using the parameter setting 'callpeak -t SAMPLE -c INPUT -f BAMPE–keep-dup = all -g mm -p 0.05 -B'. Peak summits identified by MACS2 from all samples were collected to generate a list of potential binding sites. Custom PERL script was then applied to parse the number of fragments (hereafter referred to as counts) that covered the peak summit in each sample. Each pair of properly paired-end aligned reads covering the peak summit represented one count. The total counts in each sample were normalized before comparison among samples. The potential binding sites were kept if they met all of the following conditions: (1) the sites were not located in the Mus musculus (house mouse) genome assembly GRCm38 (mm10) empirical blacklists and (2) the normalized counts in all three biological replicates in one group were larger than in its input sample, and the normalized counts in at least two replicates were more than twofold larger than in the normalized input count.

### Supplementary Figures

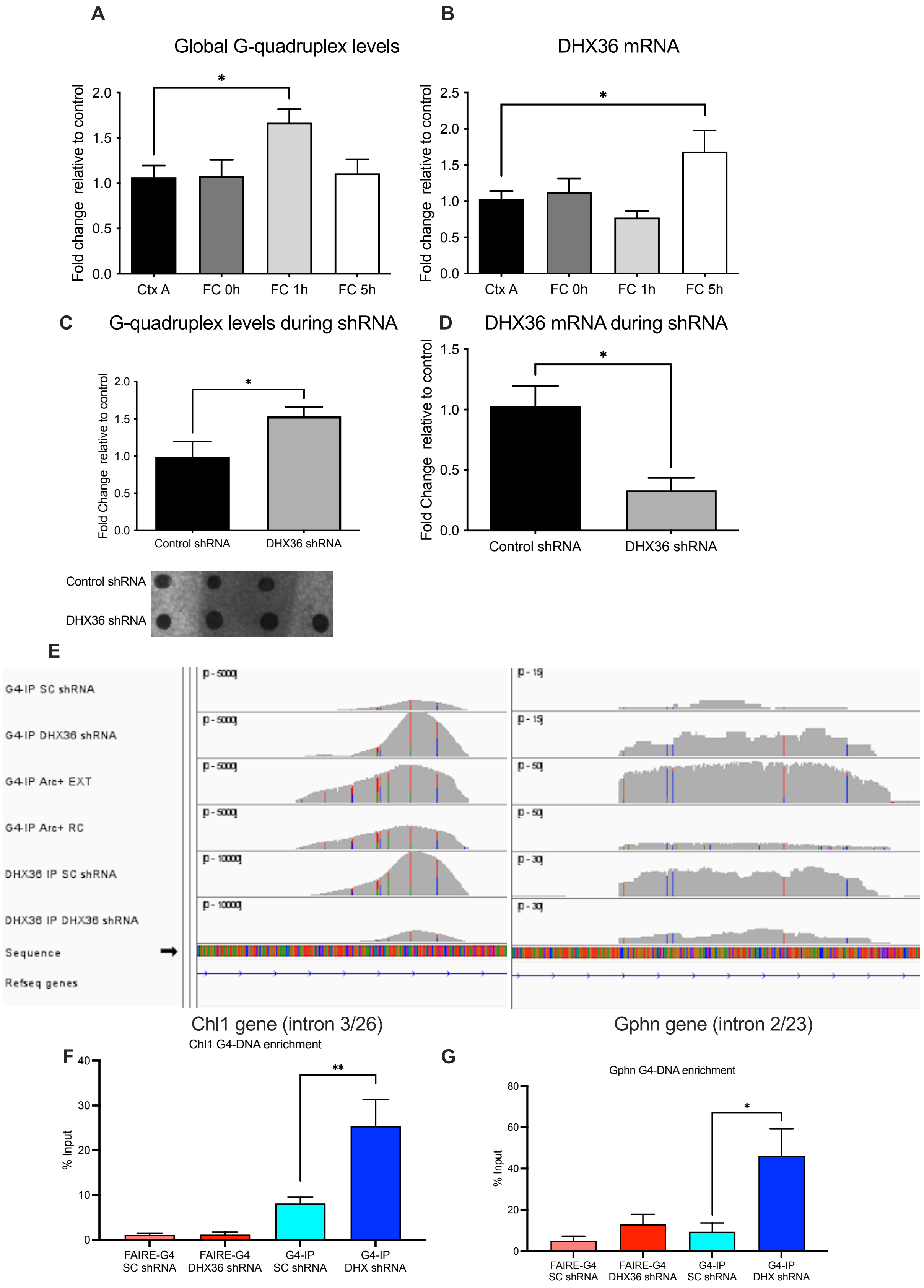


**Fig. S1. A.** In mice either exposed to context A without conditioning (Ctx A) or mice fear conditioned followed by sample collection immediately (FC 0hr), 1 hour (FC 1hr), or 5 hours post-training (FC

5hr) there was significantly more G-quadruplex at the 1hr time point compared to baseline ($F_{3,20}$ = 3.557, *p<0.05; Dunnett's post-hoc; Ctx A vs FC 0hr, p = 0.9997, Ctx A vs FC 1hr, *p = 0.0318, Ctx A vs FC 5hr p = 0.9949) **B.** There was a significant increase in the expression of the G-quadruplex specific helicase DEAH-Box Helicase 36 (DHX36) at the 5hr time point ($F_{3,18}$ = 4.631, *p<0.05; Dunnett's post-hoc; Ctx A vs FC 0hr, p = 0.9601, Ctx A vs FC 1hr, p = 0.5920, Ctx A vs FC 5hr *p = 0.0446). **C.** Application of a short hair pin RNA (shRNA) led to a significant increase in G-quadruplex DNA as measured by DNA dot blot, t(5) = 2.399, *p = 0.0308. **D.** A significant decrease in DHX36 mRNA expression, t(5) = 3.752, *p = 0.0133 **E.** Integrated genome browser (IGV) plot of G4-DNA immunoprecipitation (IP) with or without DHX36 shRNA as well as DHX36 IP with and without DHX36 shRNA **F.** Brains from SC vs DHX36 shRNA treated animals that underwent extinction training followed by G4 IP with and without reverse crosslinking were also collected. Without reverse crosslinking Formaldehyde-Assisted Isolation of Regulatory Elements (FAIRE) allows for assessment of histone and protein free regions. Assessment of the Chl1 gene revealed that much of the G4 peak had protein bound as the signal decreased, and was significantly increased following DHX36 shRNA ($F_{3,25}$ = 10.32, ***p<0.001; Dunnett's post-hoc; G4-IP SC shRNA vs. FAIRE-G4 SC shRNA, p = 0.4978, G4-IP SC shRNA vs. FAIRE-G4 DHX36 shRNA, p = 0.4531, G4-IP SC shRNA vs. G4-IP DHX shRNA, **p = 0.0098). **G.** The Gphn G4 peak appears to be mainly protein free as there was no decrease in signal in the FAIRE groups, but there was also a significant increase in G4 following DHX36 knockdown. ($F_{3,25}$ = 1.759, p>0.05; Dunnett's post-hoc; G4-IP SC shRNA vs. FAIRE-G4 SC shRNA, p = 0.9403, G4-IP SC shRNA vs. FAIRE-G4 DHX36 shRNA, p = 0.9403, G4-IP SC shRNA vs. G4-IP DHX shRNA, **p = 0.0234).

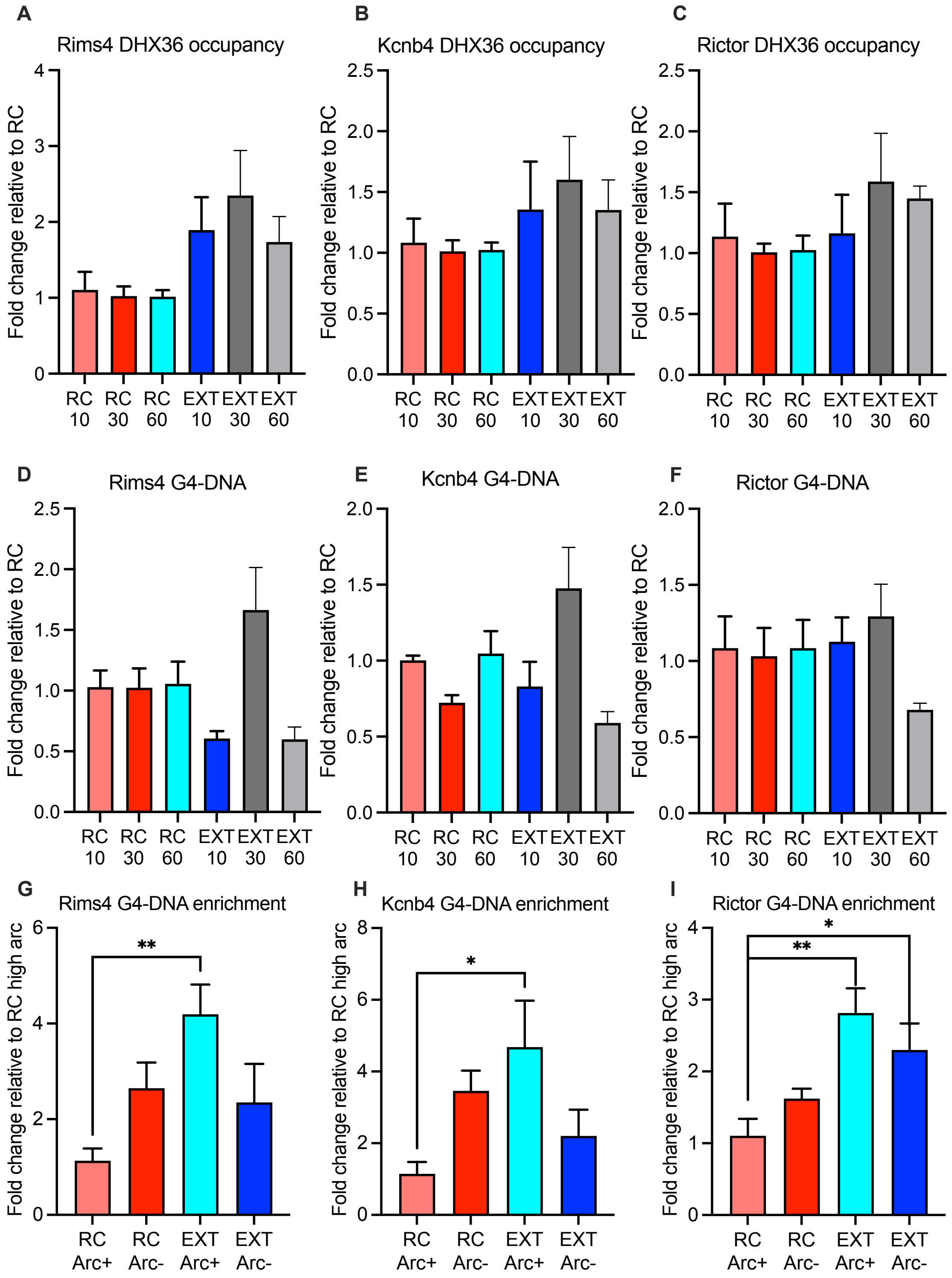


**Fig S2. DHX36 and G4 time course for increased G4-targets** DHX36 occupancy following either retention control (RC) or extinction training (EXT) for 10, 30, or 60 conditioned stimuli (10CS, 30CS, 60CS) or equivalent time for RC (RC10, RC30, RC60) with no significant differences observed for **A.** Rims4, **B.** Kcnb4, **C.** Rictor. There was also no significant change in G4 occupancy following training of the same groups and genes **D.** Rims4, **E.** Kcnb4, **F.** Rictor. Comparing G4-DNA in activated versus unsorted neurons revealed significant differences between the groups **G.** Rims4 (one-way ANOVA $F_{3,20}$= 4.157, **p<0.01; Dunnett's post-hoc; RC Arc+ vs. EXT Arc+ **p=0.0055). **H.** Kcnb4 (one-way ANOVA $F_{3,20}$= 3.511, *p<0.05; Dunnett's post-hoc; RC Arc+ vs. EXT Arc+ **p=0.01880). **I.** Rictor (one-way ANOVA $F_{3,20}$= 6.718, **p<0.01; Dunnett's post-hoc; RC Arc+ vs. EXT Arc+ **p=0.0012; RC Arc+ vs. EXT Arc- *p=0.0178).

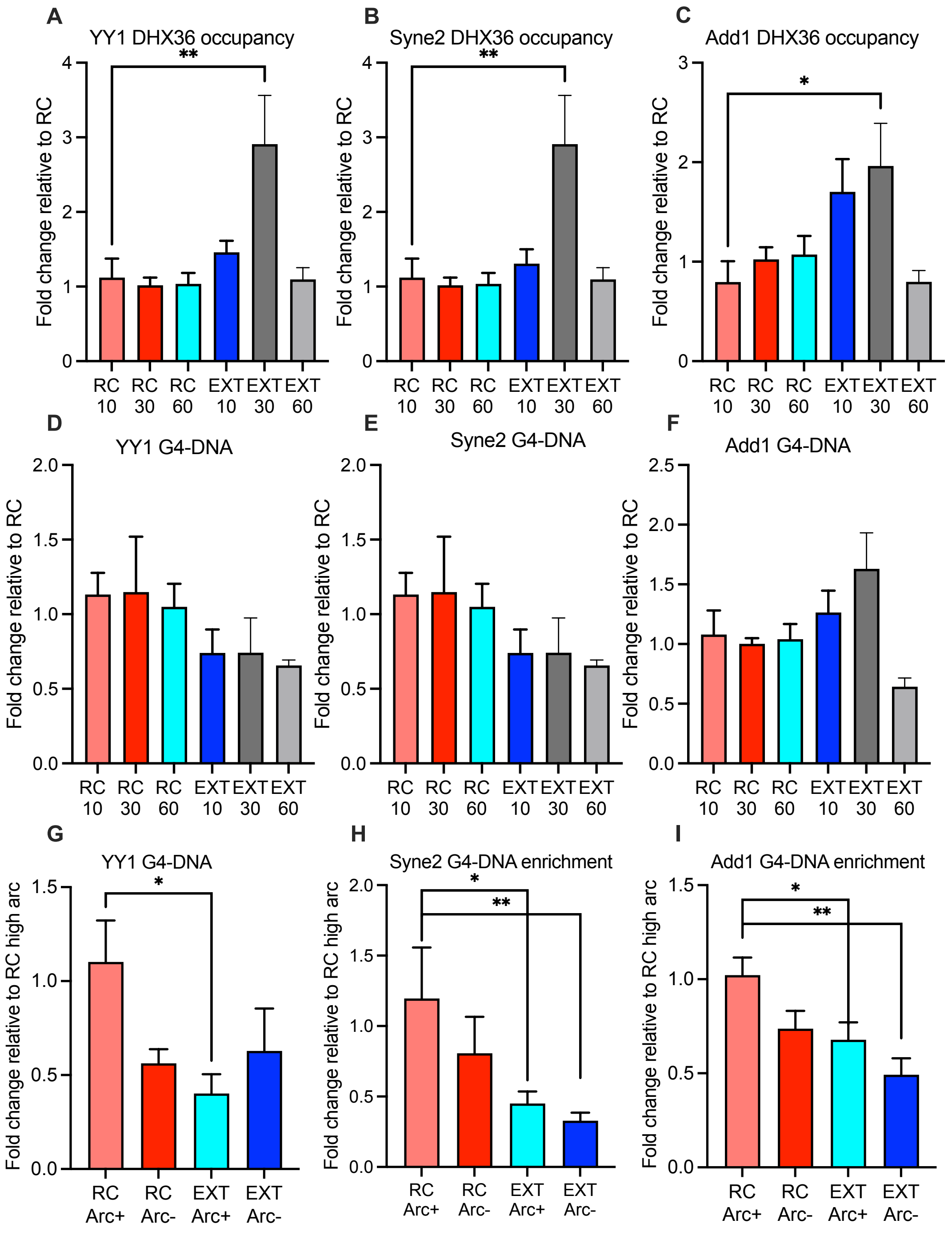


**Fig. S3. DHX36 and G4 time course for decreased G4-targets** DHX36 occupancy following either retention control (RC) or extinction training (EXT) for 10, 30, or 60 conditioned stimuli (10CS, 30CS, 60CS) or equivalent time for RC (RC10, RC30, RC60) showed a significant increase in **A.** Yy1 (one-way ANOVA $F_{5,23}$= 5.254, **p<0.01; Dunnett's post-hoc; RC10 vs EXT10 **p=0.0020). **B.** Syne2 (one-way ANOVA $F_{5,23}$= 5.304, **p<0.01; Dunnett's post-hoc; RC10 vs EXT10 **p=0.0018) and **C.** Add1 (one-way ANOVA $F_{5,23}$= 3.716, *p<0.05; Dunnett's post-hoc; RC10 vs EXT10 *p=0.0190). There was no significant change in G4 occupancy following training of the same groups and genes **D.** Yy1, **E.** Syne2, **F.** Add1. However, comparing G4-DNA in activated versus unsorted neurons revealed a significant difference between the groups for **G.** Yy1 (one-way ANOVA $F_{3,20}$= 3.382, *p<0.05; Dunnett's post-hoc; RC Arc+ vs. EXT Arc+ **p=0.0222). **H.** Syne2(one-way ANOVA $F_{3,20}$= 4.402, *p<0.05; Dunnett's post-hoc; RC Arc+ vs. EXT Arc+ *p=0.0227; RC Arc+ vs. EXT Arc- **p=0.0078). **I.** Add1 (one-way ANOVA $F_{3,20}$= 5.668, **p<0.01; Dunnett's post-hoc; RC Arc+ vs. EXT Arc+ *p=0.0401; RC Arc+ vs. EXT Arc- **p=0.0017).

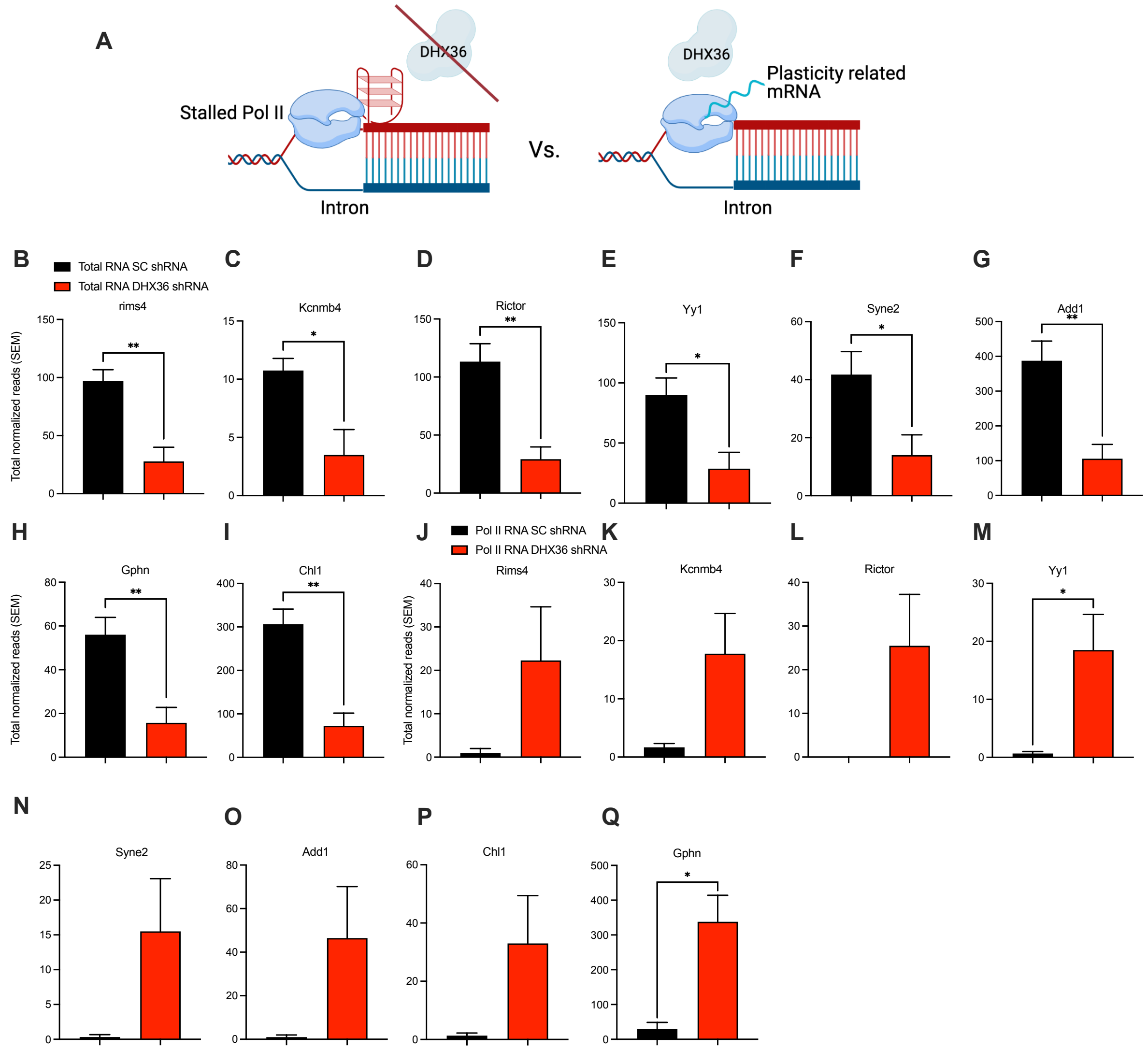


**Fig. S4. Polymerase II stalling and reduced RNA at validated targets A.** Proposed model of G4-DNA and polymerase II interaction during transcription associated with memory formation. Comparing total RNA from scrambled control (SC) and DHX36 shRNA-treated extinction-trained mice showed that there was a significant decrease in **B.** Rims4 ($t(6) = 4.416$, $^{**}p<0.01$), **C.** Kcnmb4 ($t(6) = 3.007$, $^{*}p<0.05$), **D.** Rictor ($t(6) = 4.479$, $^{**}p<0.01$), **E.** Yy1 ($t(6) = 3.135$, $^{*}p<0.05$), **F.** Syne2 ($t(6) = 2.618$, $^{*}p<0.05$), **G.** Add1($t(6) = 4.026$, $^{**}p<0.01$), **H.** Chl1 ($t(6) = 5.175$, $^{**}p<0.01$), **I.** Gphn ($t(6) = 3.797$, $^{**}p<0.01$). Pol II bound RNA increased in **J.** Rims4 ($t(5) = 1.448$, $p=0.1036$), **K.** Kcnmb4 ($t(5) = 1.906$, $p=0.0537$), **L.** Rictor ($t(5) = 1.833$, $p=0.0631$), **M.** Yy1 ($t(5) = 2.451$, $^{*}p<0.05$), **N.** Syne2 ($t(5) = 1.691$, $p=0.0758$) **O.** Add1 ($t(5) = 1.629$, $p=0.0821$) **P.** Chl1 ($t(5) = 1.630$, $p=0.0820$) **Q.** Gphn ($t(5) = 3.351$, $^{*}p<0.05$)

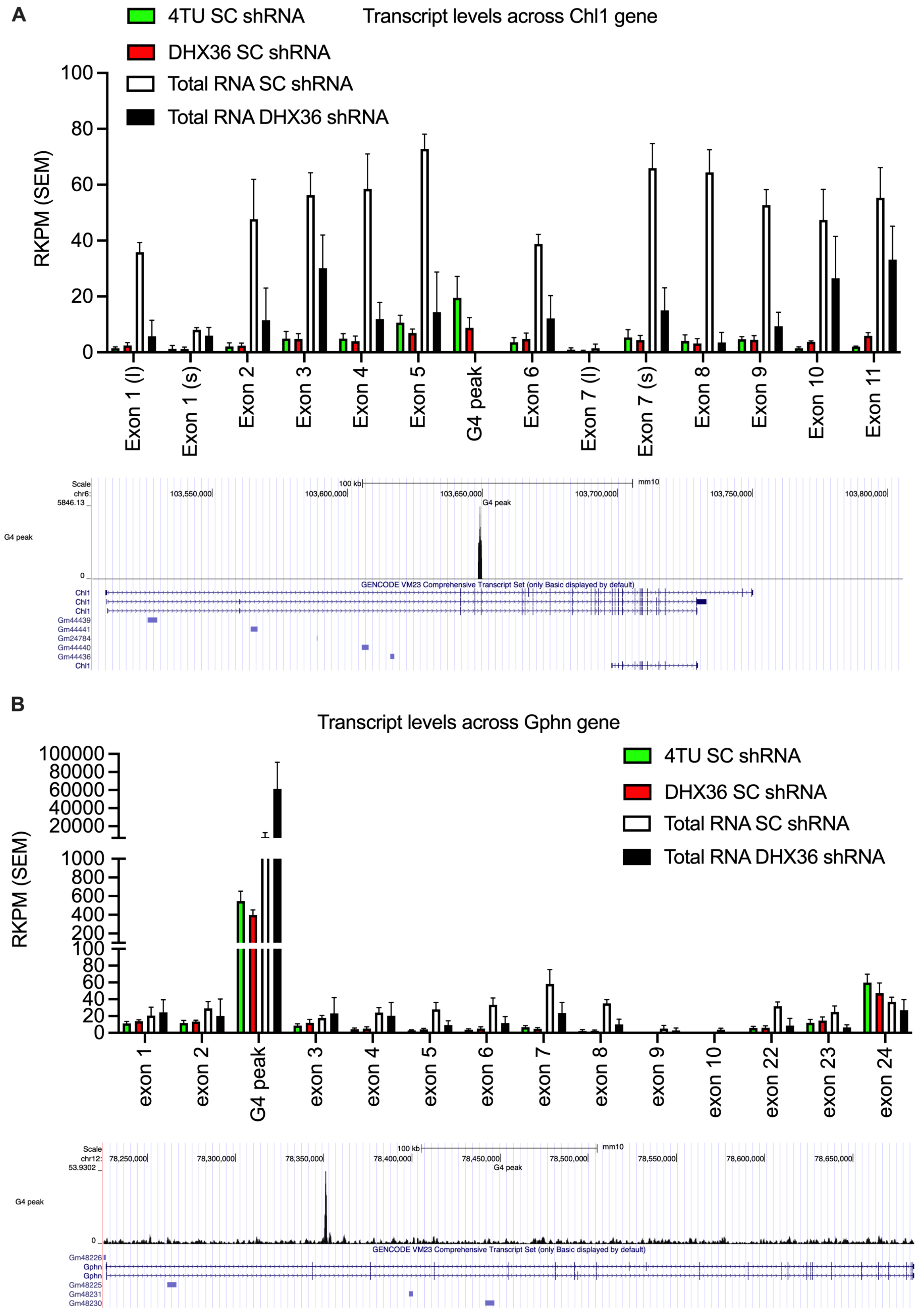


**Fig. S5. Chl1 and Gphn total and nacent RNA overlap with G4-sites** Integrated genome browser (IGV) showing the reads per kilobase million (RPKM) derived from total RNA sequencing (total RNA) and 4-thiouracil (4TU) metabolic labelling sequencing of mice treated with either scrambled control or DHX36 shRNA. Data is across each exon of the gene relative to the G4-peak for: **A.** Chl1 and **B**. Gphn.

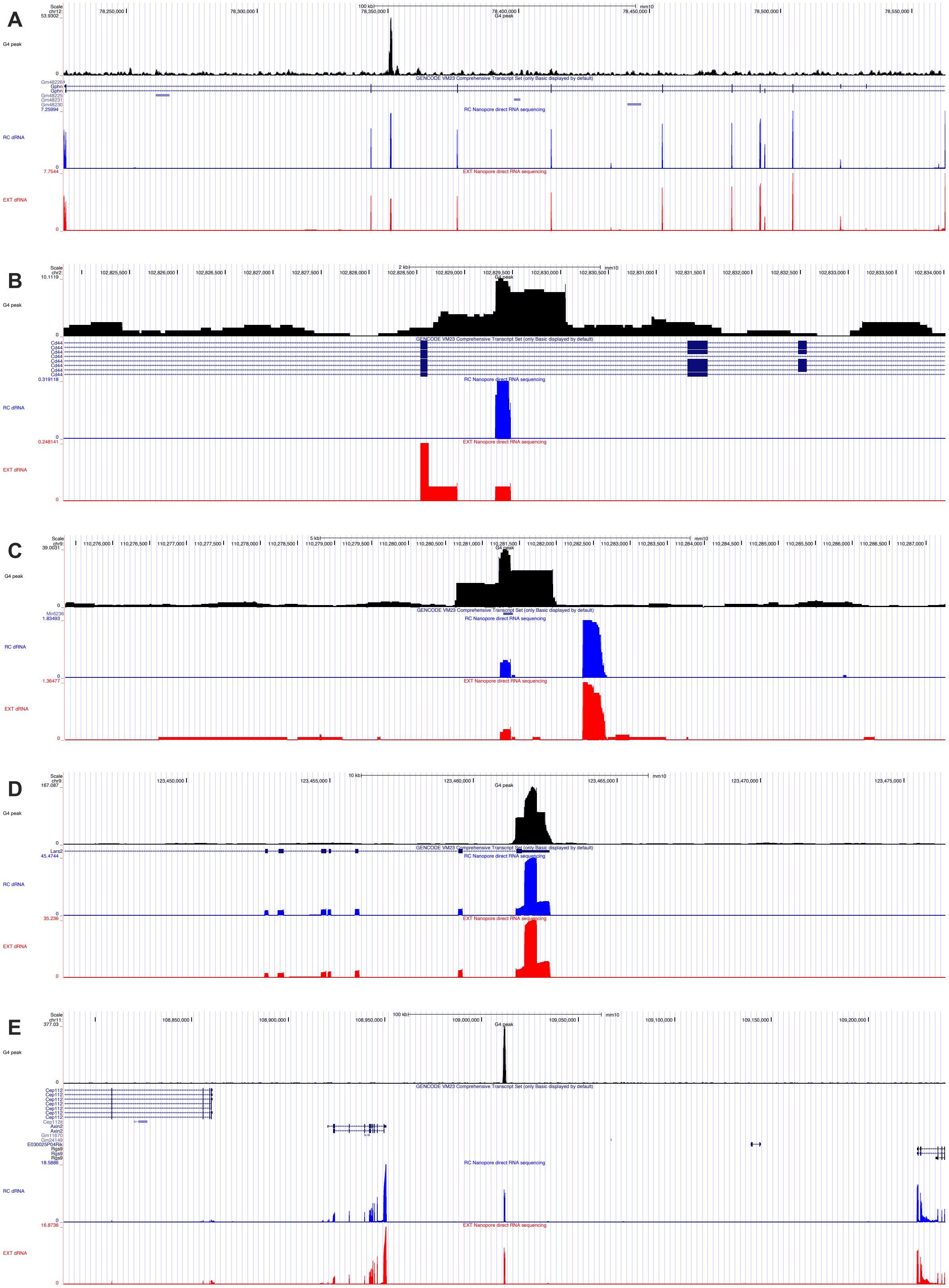

A
Scale
chr12:
G4 peak
GENCODE VM23 Comprehensive Transcript Set (only Basic displayed by default)
RC Nanopore direct RNA sequencing
RC dRNA
EXT Nanopore direct RNA sequencing
EXT dRNA
B
Scale
chr2:
G4 peak
Cd44
RC dRNA
EXT dRNA
C
Scale
chr9:
G4 peak
RC dRNA
EXT dRNA
D
Scale
chr9:
G4 peak
Lars2
RC dRNA
EXT dRNA
E
Scale
chr11:
G4 peak
Cep112
Axin2
Rgs9
RC dRNA
EXT dRNA

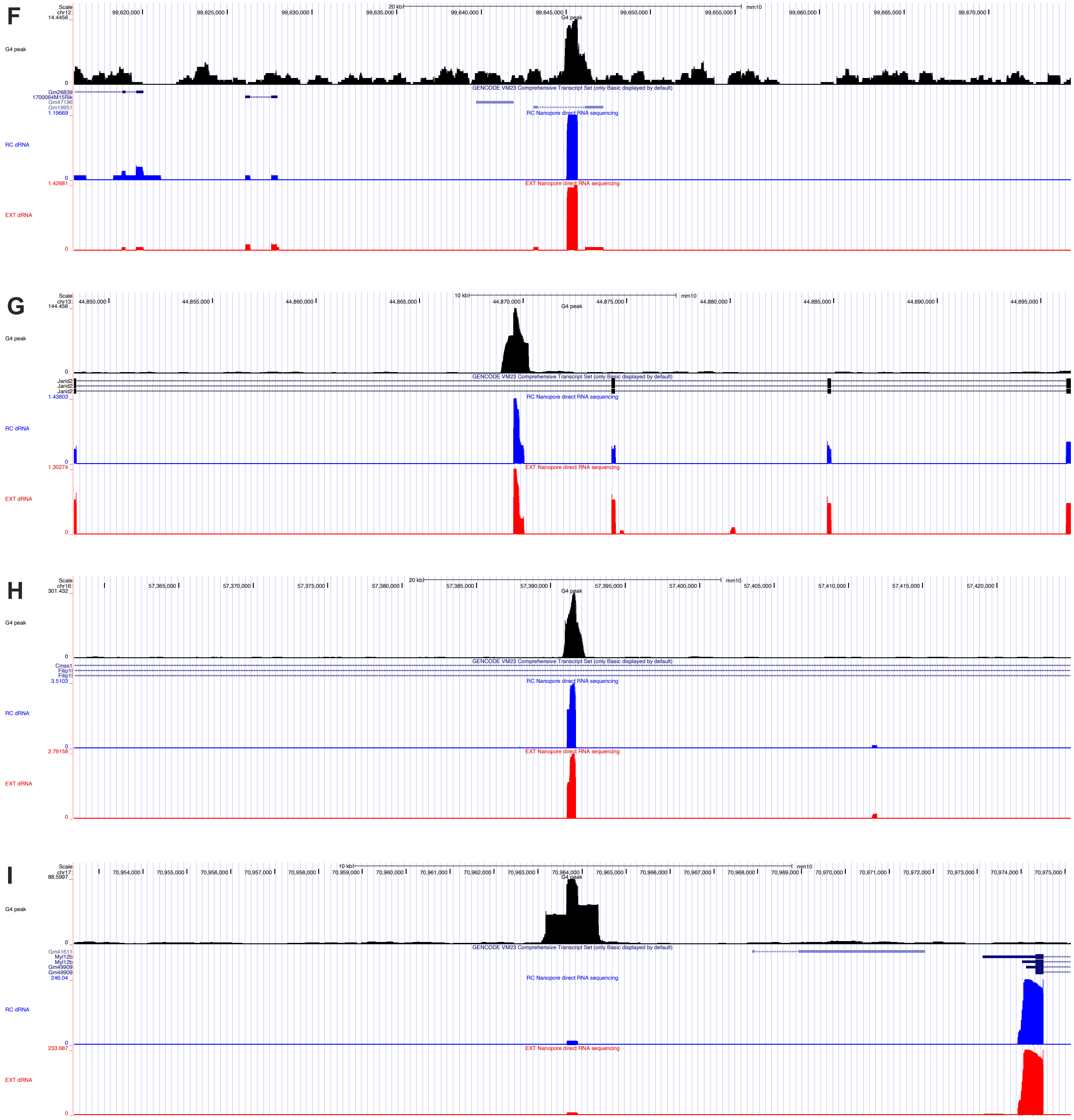


**Fig. S6. Sites of G4-DNA and significant RNA change across the transcriptome** Integrated genome browser (IGV) traces of the G-quadruplex peak, as well as RNA expression across a variety of genomic loci including **A.** Gphn, **B.** Cd44, **C.** Mir6236 **D.** Lars2, **E.** Intragenic site 1, **F.** Gm19951, **G.** Jarid2, **H.** Cmss1, **I.** Intragenic site 2

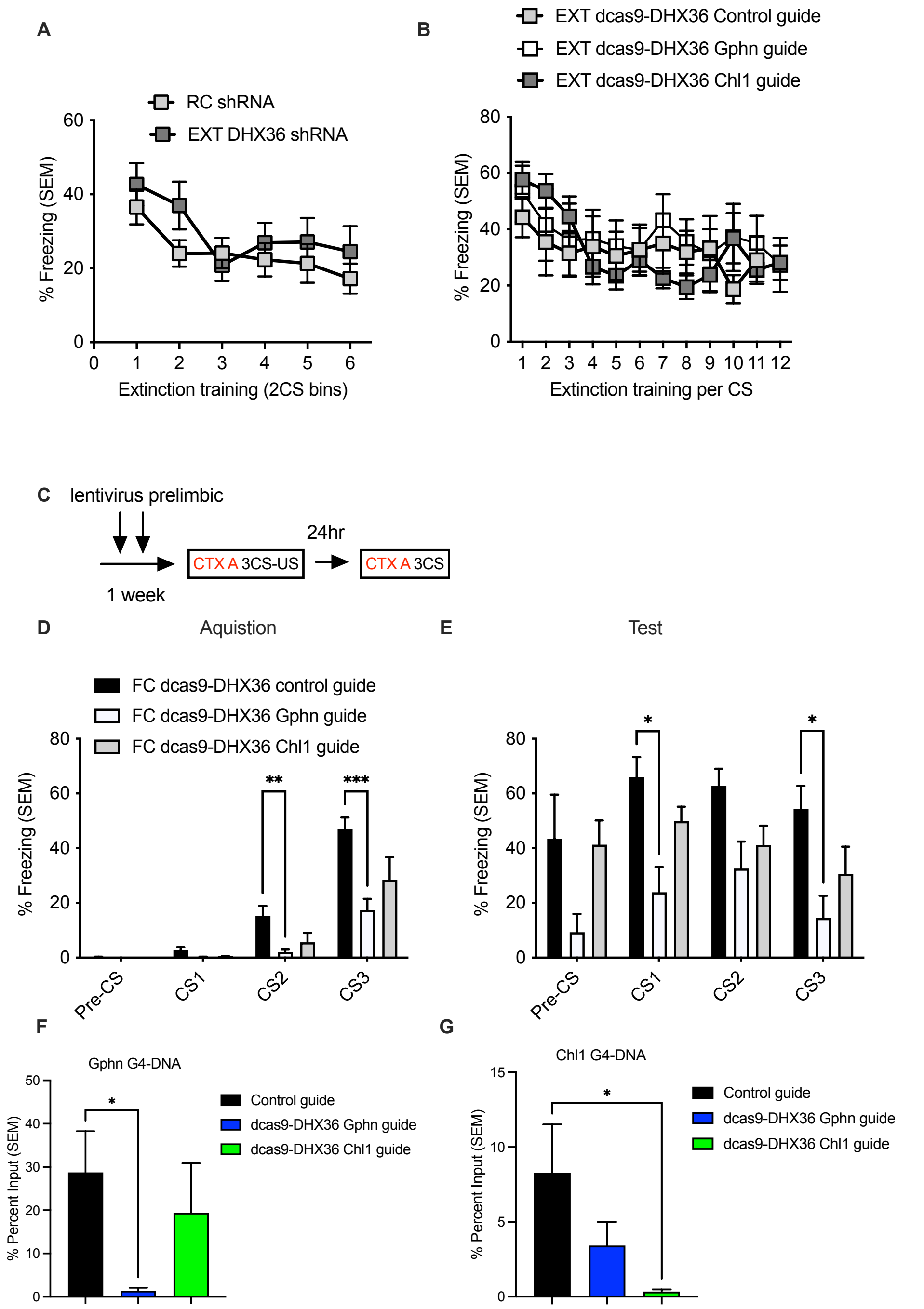


**Fig. S7. Fear and extinction behaviour during viral manipulation A.** During extinction training of SC and DHX36 shRNA infused animals there was no significant difference in freezing **B.** Similarly, during extinction training of dcas9-DHX36 infused animals there was no significant difference in freezing **C.** Pictorial of timeline for behavioural training and lentivirus infusion **D.** After infusion of dcas9-DHX36 there was a significant impairment in acquisition for the Gphn guide ($F_{2,30} = 6.377$, $**p<0.01$; Dunnett's post-hoc; CS1 Control guide vs. Gphn guide p = 0.0573, CS2 Control guide vs. Gphn guide **p = 0.0040 CS3 Control guide vs. Gphn guide ***p = 0.0002). **E.** There was also a significant impairment in recall for the Gphn guide **F.** Following infusion of dcas9-DHX36 with either control guides, Gphn guides, or Chl1 guides there was a significant reduction in G4-DNA at the Gphn locus following Gphn guide infusion ($F_{2,10} = 15.65$, $***p<0.001$; Dunnett's post-hoc; CS1 Control guide vs. Gphn guide *p = 0.0211, CS2 Control guide vs. Gphn guide p = 0.0696 CS3 Control guide vs. Gphn guide *p = 0.0301). **G.** In a separate cohort of animals there was a significant reduction in G4-DNA at the Chl1 locus with the Chl1 guides ($F_{2,24} = 3.517$, $*p<0.05$; Dunnett's post-hoc; Control guide vs. Gphn guide p =0.0906, Control guide vs Chl1 guide *p = 0.0290).

**Data S1. (separate file)**
G4 and DHX36 immunoprecipitation sequencing for scrambled control and DHX36 shRNA treated extinction trained mice

**Data S2. (separate file)**
FACS sorted G4 sequencing for scrambled control and DHX36 shRNA treated extinction trained mice

**Data S3. (separate file)**
Total RNA counts for scrambled control and DHX36 shRNA treated extinction trained mice

**Data S4. (separate file)**
4-thiouracil counts for scrambled control and DHX36 shRNA treated extinction trained mice

**Data S5. (separate file)**
Counts for polymerase II associated RNA from scrambled control and DHX36 shRNA treated extinction trained mice

**Data S6. (separate file)**
Classification of location of all significant total and 4-thiouracil RNA for scrambled control and DHX36 shRNA treated extinction trained mice

**Data S7. (separate file)**
Comparison of total and 4-thiouracil RNA for scrambled control and DHX36 shRNA treated extinction trained mice

**Data S8. (separate file)**
Table of primers, antibodies and reagents required for experiments
